## Supplementary Figures for "Coral larvae employ nitrogen sequestration mechanisms to stabilize carbon provisioning from algal symbionts under increased temperature"

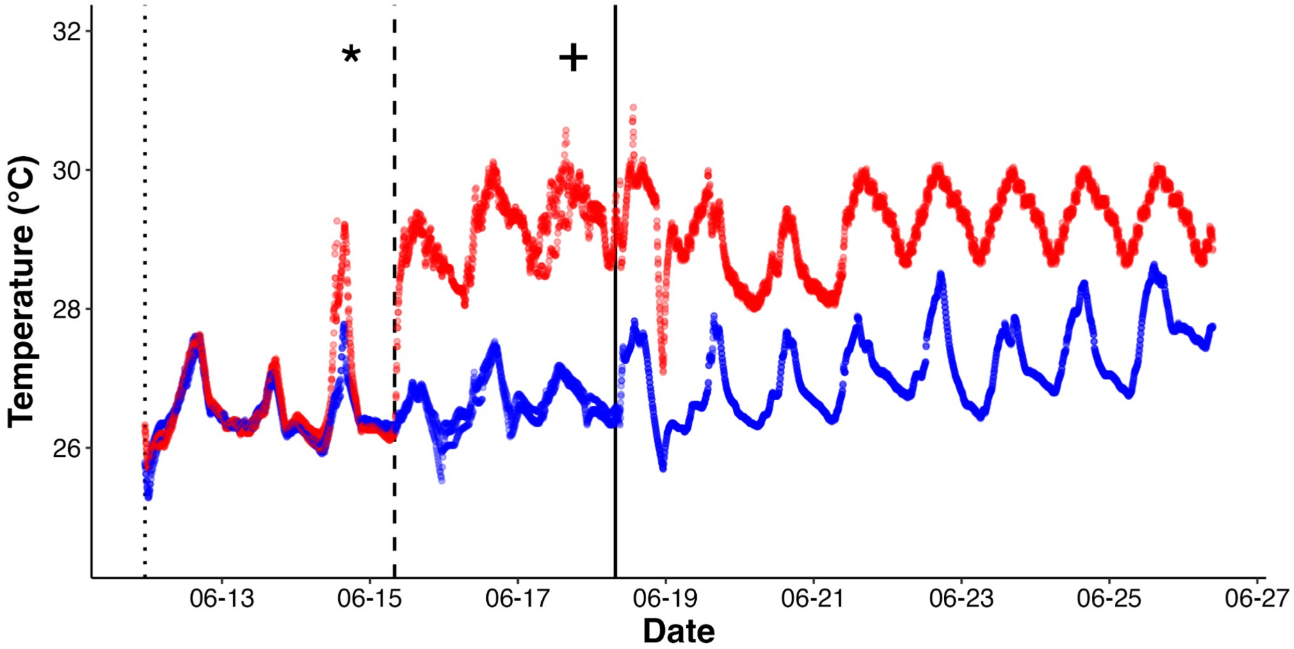


**Fig S1.** Temperature treatments (ambient = blue; high = red) during each period of the study: embryonic development (dotted line; 06-12 through 06-14), larval exposure (dashed line; 06-15 through 06-18), and settlement/post-settlement survival (solid line; 06-19 through 06-26). X-axis indicates date (MM-DD). Asterisks (*) indicate the start of temperature treatments (06-14) and plus (+) indicates time of larval sampling (06-18). Temperature recorded every 15 min by n=3 loggers per temperature treatment.

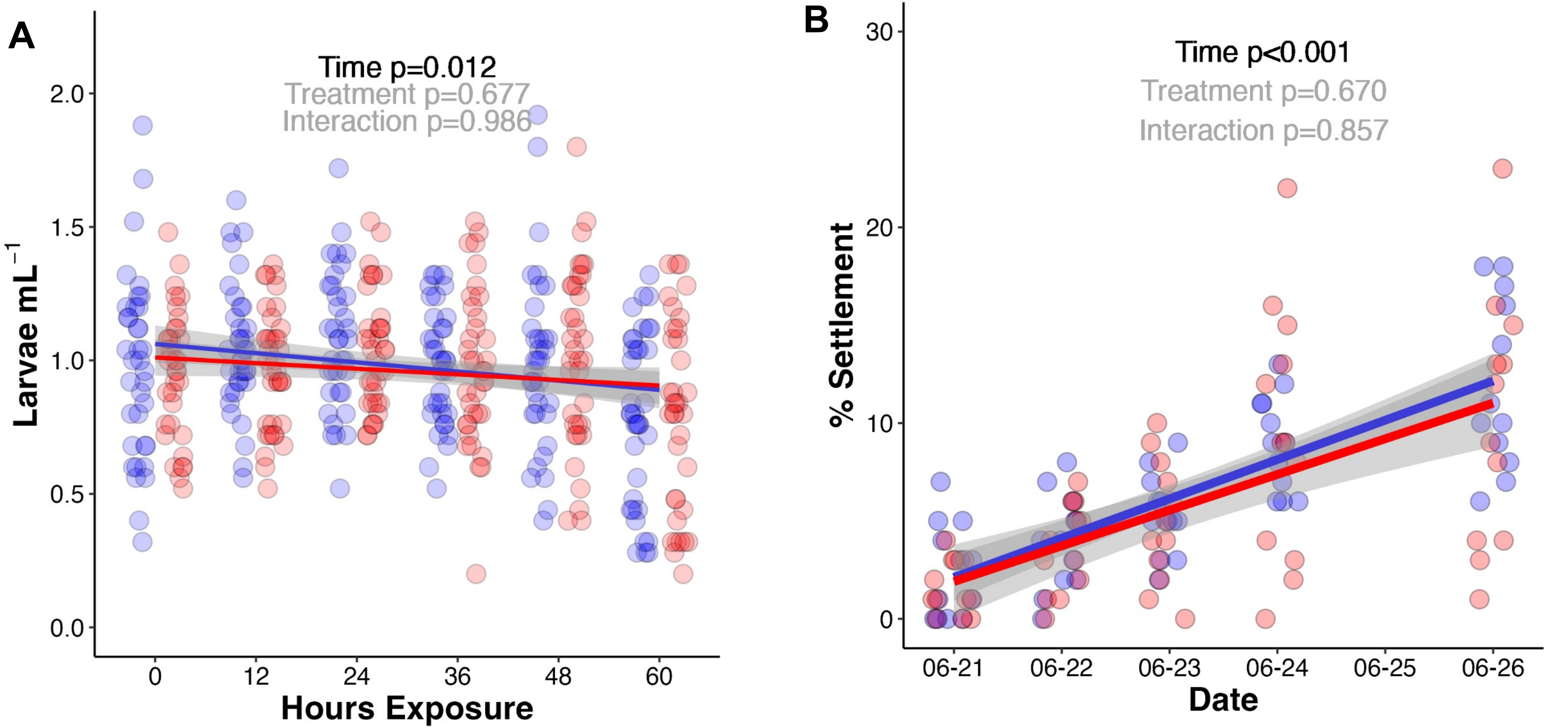


**Fig S2.** *Montipora capitata* larval (A) survival during 3 days of exposure to ambient (blue) and high (red) temperature treatments. (B) Percent larval settlement following larval exposure. Following larval exposure, larvae were settled under temperature treatments and held at these treatments over a 5-day period. Effects of time and rearing treatment were tested using linear mixed effect models with tank as a random intercept (black $p<0.05$; gray $p>0.05$). Linear model fit line shown with gray indicating 95% confidence intervals.

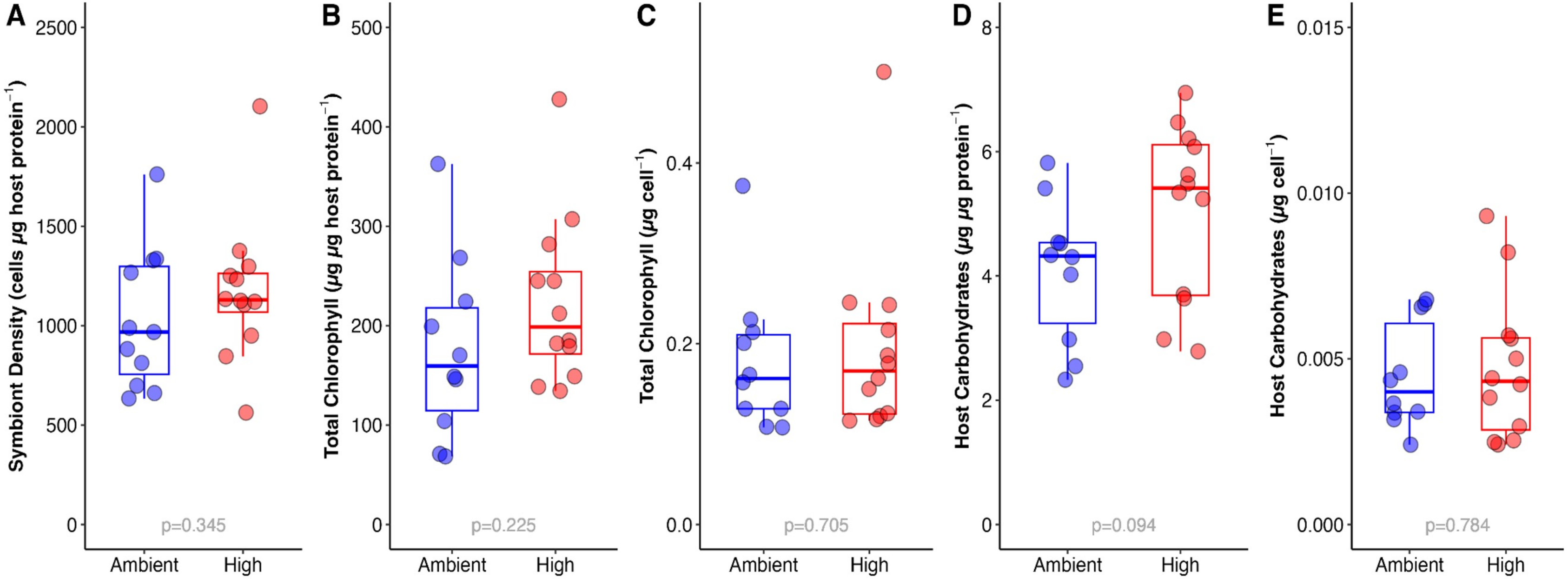


**Fig S3.** *Montipora capitata* larval physiological measurements following 3 days of exposure to ambient (blue) and high (red) temperature treatments. (A) Symbiont cell density, cells per μg host protein content; (B) Symbiont chlorophyll (α + c2), μg per μg host protein content; (C) Symbiont chlorophyll (α + c2), μg symbiont cell; (D) Host carbohydrate content, μg per μg host protein content; (E) Host carbohydrate content, μg per symbiont cell; Effect of treatment was tested using Welch t-tests (black $p<0.05$; gray $p>0.05$).

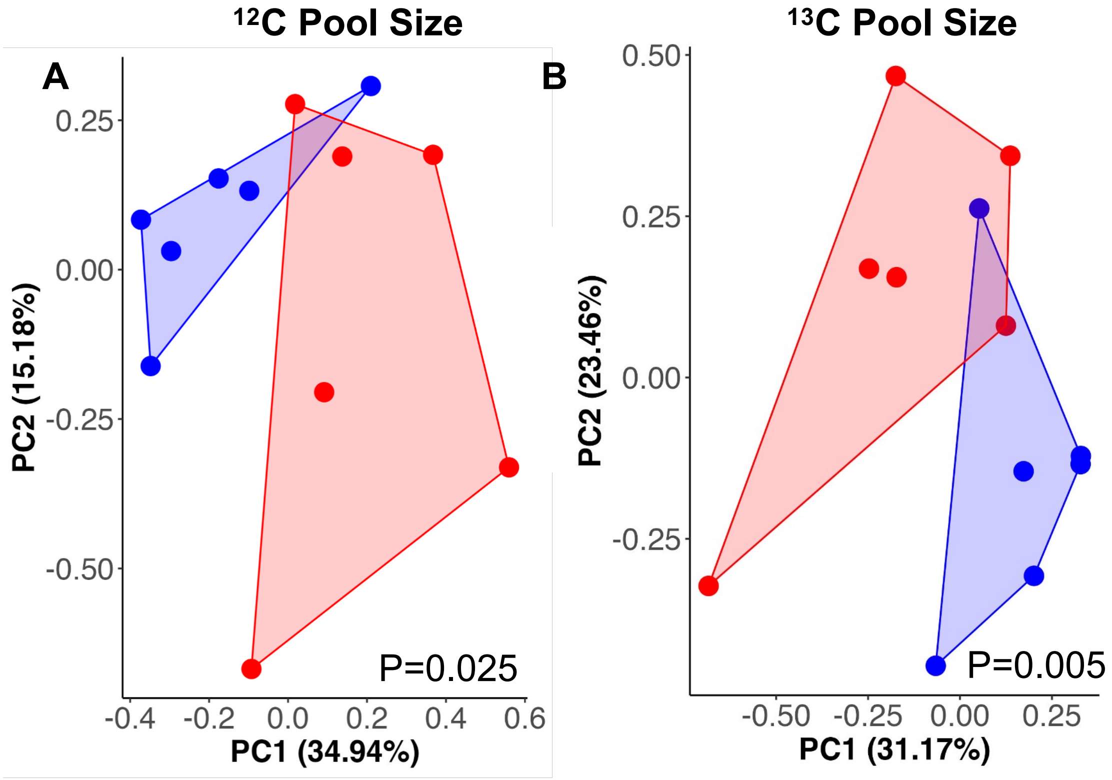


**Fig S4.** Multivariate visualization of metabolite pool size in *Montipora capitata* larval samples. (A) Principal components analysis of metabolite pool sizes in larvae incubated with unlabeled ($^{12}$C) sodium bicarbonate between temperature treatments. (B) Principal components analysis of metabolite pool sizes in larvae incubated with labeled ($^{13}$C) sodium bicarbonate between temperature treatments. Blue indicates larvae exposed to ambient temperature; red indicates larvae exposed to high temperature. Axis show percent variance explained by each principal component.  P-values indicate significance of temperature treatment on multivariate pool size analyzed using PERMANOVA analyses.

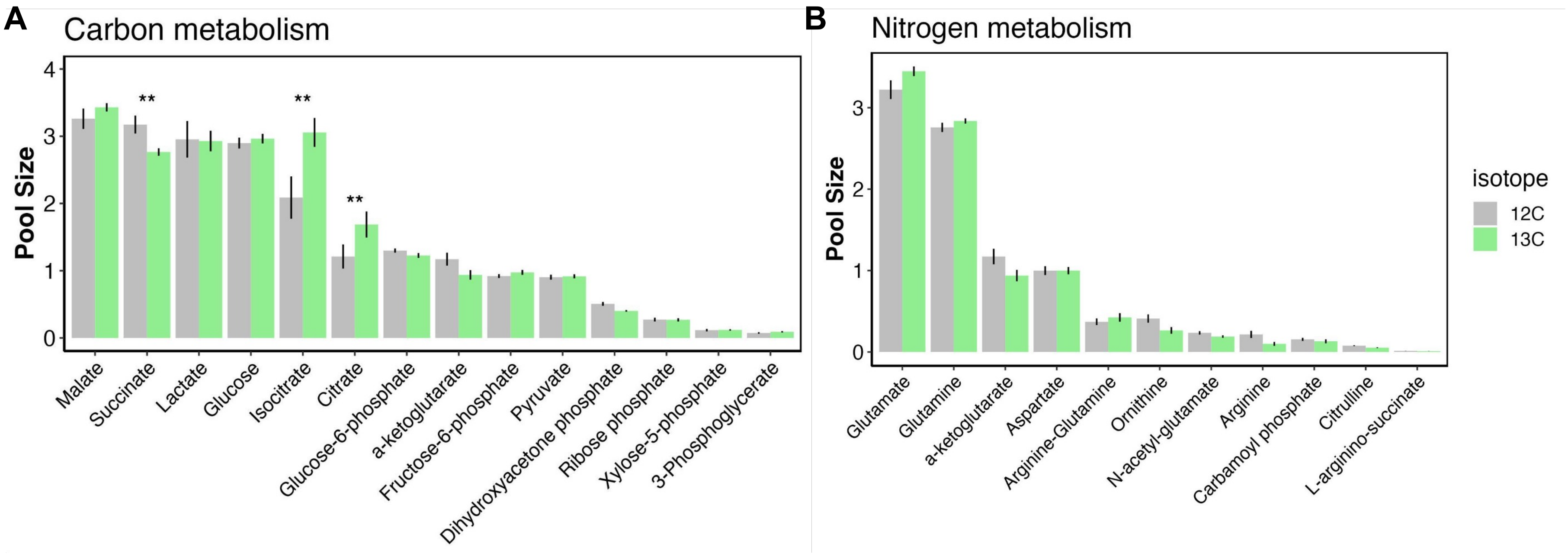


**Fig S5. Pool size of metabolites between isotope treatments.** Mean (± standard error of mean) pool size in metabolites of interest related to (A) carbon metabolism, including glycolysis, pentose phosphate pathway, and the tricarboxylic acid cycle, and (B) nitrogen metabolism, including ammonium assimilation and the urea cycle in $^{12}C$ (gray) and $^{13}C$ (green) isotope treatments. Error bars represent standard error of mean. ** indicates $p<0.01$ as determined by estimated marginal means posthoc tests.

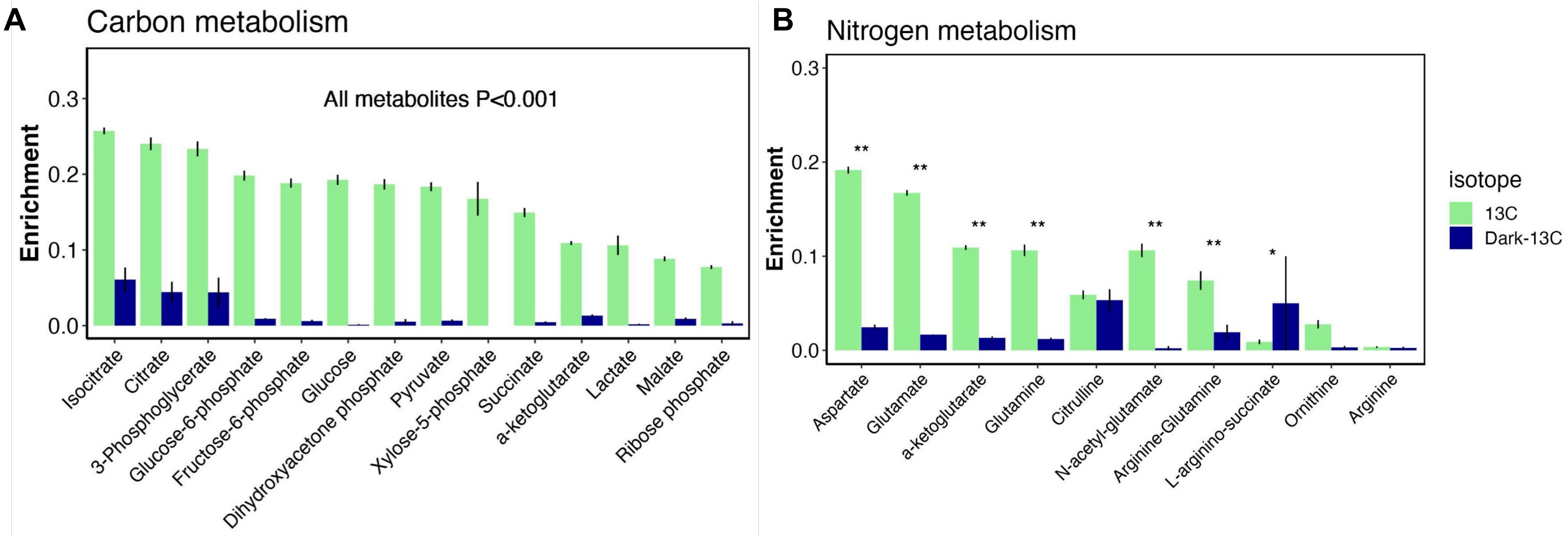


**Fig S6. Enrichment in metabolites of interest between $^{13}C$ light and dark treatments.** Enrichment in metabolites of interest related to (A) carbon and (B) nitrogen metabolism in larvae incubated with unlabeled $^{12}C$ sodium bicarbonate (gray), and labeled $^{13}C$ sodium bicarbonate (green). Error bars represent standard error of mean. * indicates p<0.05; ** indicates p<0.01 as determined by estimated marginal means posthoc tests. In (A), all metabolites were significant at P<0.001.

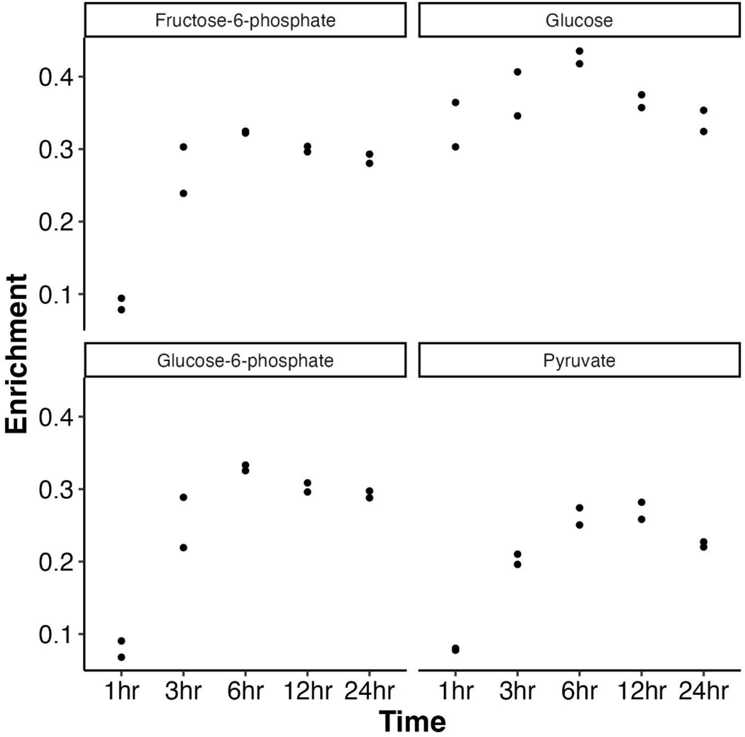


**Fig S7.** *Montipora capitata* larval stable isotope ($^{13}C$ sodium bicarbonate; 4 mM) metabolomic time series methodological control. Isotopic label enrichment shown for for glucose (a primary photosynthate), glycolysis intermediates (fructose-6-phosphate, glucose-6-phosphate), and a glycolysis end product (pyruvate) across a 24 hr time series sampling.

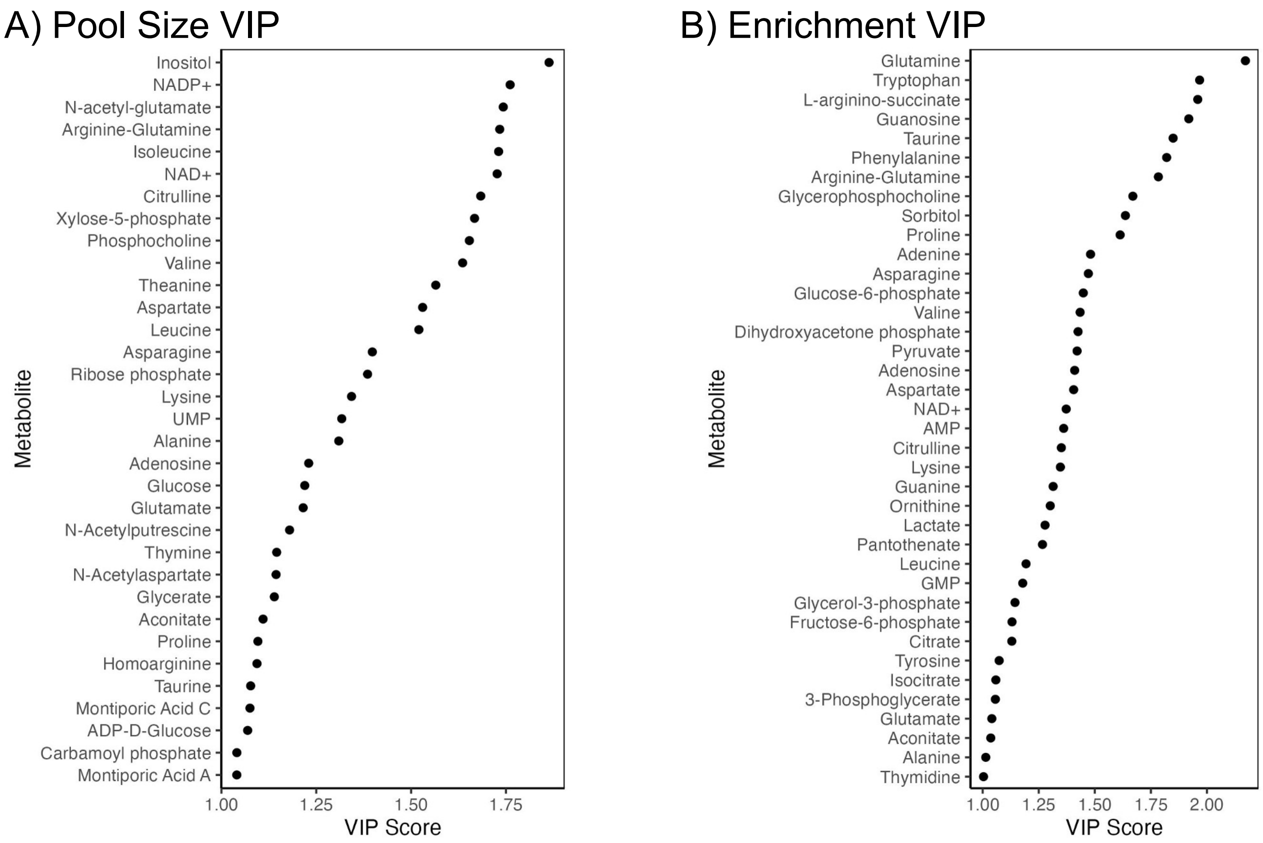


**Fig S8. Variable Importance in Prediction.** Metabolite Variable Importance in Prediction (VIP) scores for metabolites with VIP scores > 1 that significantly drive differences in (A) pool size and (B) $^{13}C$ enrichment between high and ambient temperature metabolomic responses in *Montipora capitata* larvae. VIPs were identified through partial least squares discriminant analysis (PLS-DA).

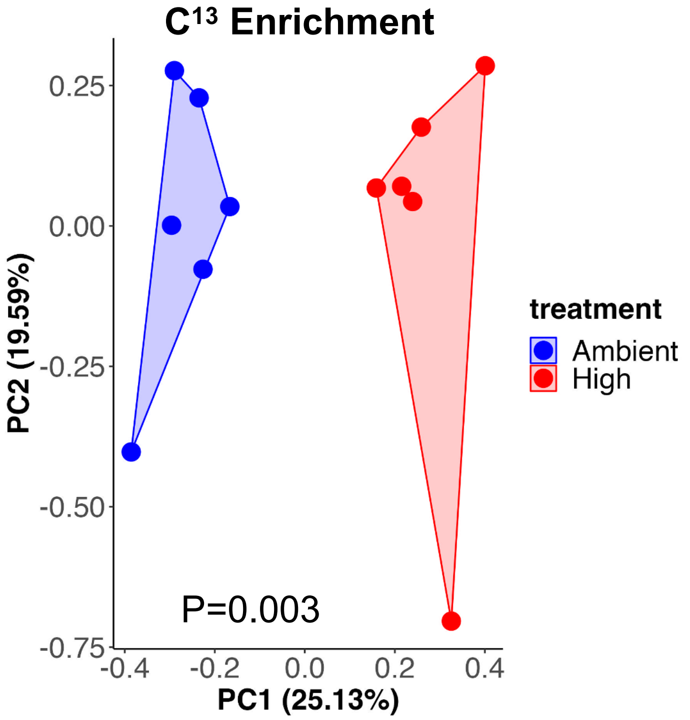


**Fig S9.** Multivariate visualization of metabolite $^{13}C$ enrichment in *Montipora capitata* larvae exposed to ambient (blue) and high (red) temperatures. Axes show percent variance explained by each principal component. P-value indicates significance of temperature treatment on multivariate enrichment analyzed using PERMANOVA analyses.

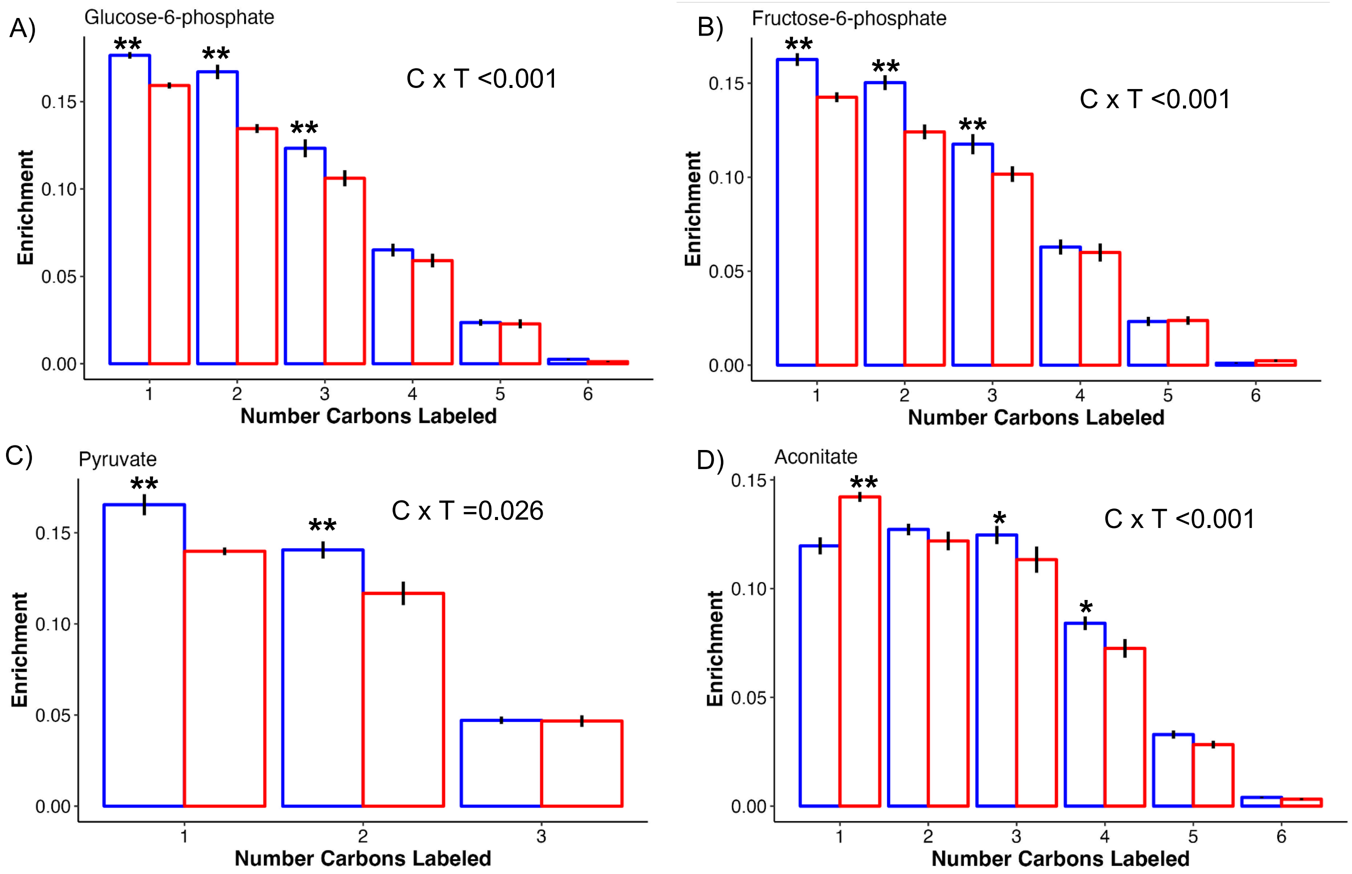


**Fig S10. Metabolite carbon specific labeling.** Metabolite enrichment for molecules of each metabolite with the specified number of labeled carbons. Significance of the interaction of number of labeled carbons (C) and temperature treatment (T) shown in text. In all plots, red indicates high temperature and blue indicates ambient. Error bars represent standard error of mean. Asterisks indicate significance of posthoc comparisons with * indicating $p<0.05$ and ** indicating $p<0.01$.
